# TOR represses stress responses through global regulation of H3K27 trimethylation in plants

**DOI:** 10.1101/2021.03.28.437410

**Authors:** Yihan Dong, Veli V. Uslu, Alexandre Berr, Gaurav Singh, Csaba Papdi, Victor A Steffens, Thierry Heitz, Lyubov Ryabova

## Abstract

Target of Rapamycin (TOR) functions as a central sensory hub to link a wide range of external stimuli to gene expression. However, the mechanisms underlying stimulus-specific transcriptional reprogramming by TOR remains elusive. Our *in silico* analysis in Arabidopsis demonstrates that TOR-repressed genes are associated with either bistable or silent chromatin states. Both states regulated by TOR signaling pathway are associated with high level of H3K27me3 deposited by CURLY LEAF (CLF) in specific context with LIKE HETEROCHROMATIN PROTEIN1 (LHP1). Combinations of epigenetic modifications H3K4me3 and H3K27me3 implicate bistable feature which alternates between on and off state allowing rapid transcriptional changes upon external stimuli. Chromatin remodeler SWI2/SNF2 ATPase BRAHMA (BRM) activates TOR-repressed genes only at bistable chromatin domains to rapidly induce biotic stress responses. Here we demonstrated both *in silico* and *in vivo* that TOR represses transcriptional stress responses through global maintenance of H3K27me3.

## Introduction

The protein kinase TOR is a conserved sensor of nutrient availability and energy status in eukaryotes, known to regulate many fundamental cellular processes, such as translation, autophagy and cell cycle (1). In animals, since diet and nutrient exert transgenerational effect, correlations between TOR functions and epigenome started to be explored (2). Besides the known relationship between histone acetylation and mammalian TOR (mTOR) signaling pathway, mammalian/mechanistic TOR complex 1 (mTORC1) was recently found to activate its canonical substrate S6K1 which in turn phosphorylates histone H2B to influence histone methylation(2). Unlike in animals, little is known in plants to link TOR function to epigenetic modifications.

As sessile organisms, plants continuously and rapidly adjust their development to a multitude of environmental stresses. Beyond stress sensing and signal transduction, plant TOR tunes developmental plasticity via its capacity to re-program the transcriptome (3). This global transcriptional reprogramming is partially mediated by the retention of ethylene-insensitive protein 2 (EIN2) in the cytoplasm to allow gene expression involved in DNA replication and cell wall biosynthesis (4). However, it still remains obscure how an active TOR can suppress stress response and defense mechanism at transcriptional level. Among other mechanisms, epigenetic modifications such as DNA methylation and histone modifications have emerged as fundamental mediators in controlling gene expression.

Multiple combinations of different epigenetic modifications exist at the whole genome level and contribute to the complexity of epigenetic landscapes. By reducing their dimensionality, a detailed bioinformatic integration of epigenomic data defines distinctive chromatin states (CS) in *A. thaliana* (5). The two silent states CS-8 and 9 are enriched in H3K9me2 and H3K27me1 and preferentially mark constitutive heterochromatin. Another inactive state CS-5 represents the typical Polycomb-regulated facultative heterochromatin with abundant H3K27me3. Meanwhile, CS-1, 3 and 6 are characterized by high amounts of active marks (*e.g*., H3K4me3, H3 acetylation, H3K36me3) and typically found at actively transcribed genes. Despite their opposite effect on transcription, H3K4me3 and H3K27me3 can co-reside at a number of dynamically regulated genes in both animals and plants. This feature is the hallmark of CS-2, a state preferentially found at the promoter of genes with transcript amounts similar to the ones marked by CS-5. A bistable model of regulation achieved through the delicate balance between H3K4me3 and H3K27me3 was proposed to poise silenced genes marked by CS-2 for rapid activation upon need (6, 7). Adding additional layers of complexity, chromatin-binding proteins like H3K27me3 reader LIKE HETEROCHROMATIN PROTEIN 1 (LHP1) or SWI2/SNF2 ATPase BRAHMA (BRM) further subdivide these chromatin domains into distinct subdomains (8–10).

In this study, we show that specific epigenetic features represent a dynamic hallmark of stimulus-specific transcriptional responses mediated by TOR signaling upon environmental stimuli. We demonstrate that active TOR represses genes associated with bistable (CS2) or silent chromatin domains (CS5) by mediating the deposition of H3K27me^3^ via CLF and LHP1. We propose a model in which environmental stimuli inhibit TOR, which subsequently allows BRM to reduce repression of bistable domains to rapidly activate biotic stress specific gene expression.

## Results

### TOR-repressed genes are associated with CS-2 and CS-5 at the transcription start site

To assess whether specific functional categories are enriched in any of the previously described chromatin states, we performed gene ontology (GO) enrichment analysis (5). Arabidopsis genes were individually assigned to one of the nine chromatin states based on the chromatin state present at their TSS. Surprisingly, each of the nine chromatin states were enriched in distinct functional categories (Fig. 1A and Table S1). Indeed, genes related to protein and RNA processes appeared highly enriched in CS-1, 3 and 6, which are characterized by the presence of active epigenetic marks. The Polycomb-associated CS-5 contained significantly more transcription, cell differentiation, and redox-regulation related genes. Abiotic and biotic stress genes are overrepresented in the bistable CS-2.Worth noting, functional categories enriched in CS-2 resemble the primary target genes in stress and immune response pathways repressed by TOR (11). Because of this similarity, we next search for particular chromatin state enrichments among TOR repressed genes.

**Fig. 1:**
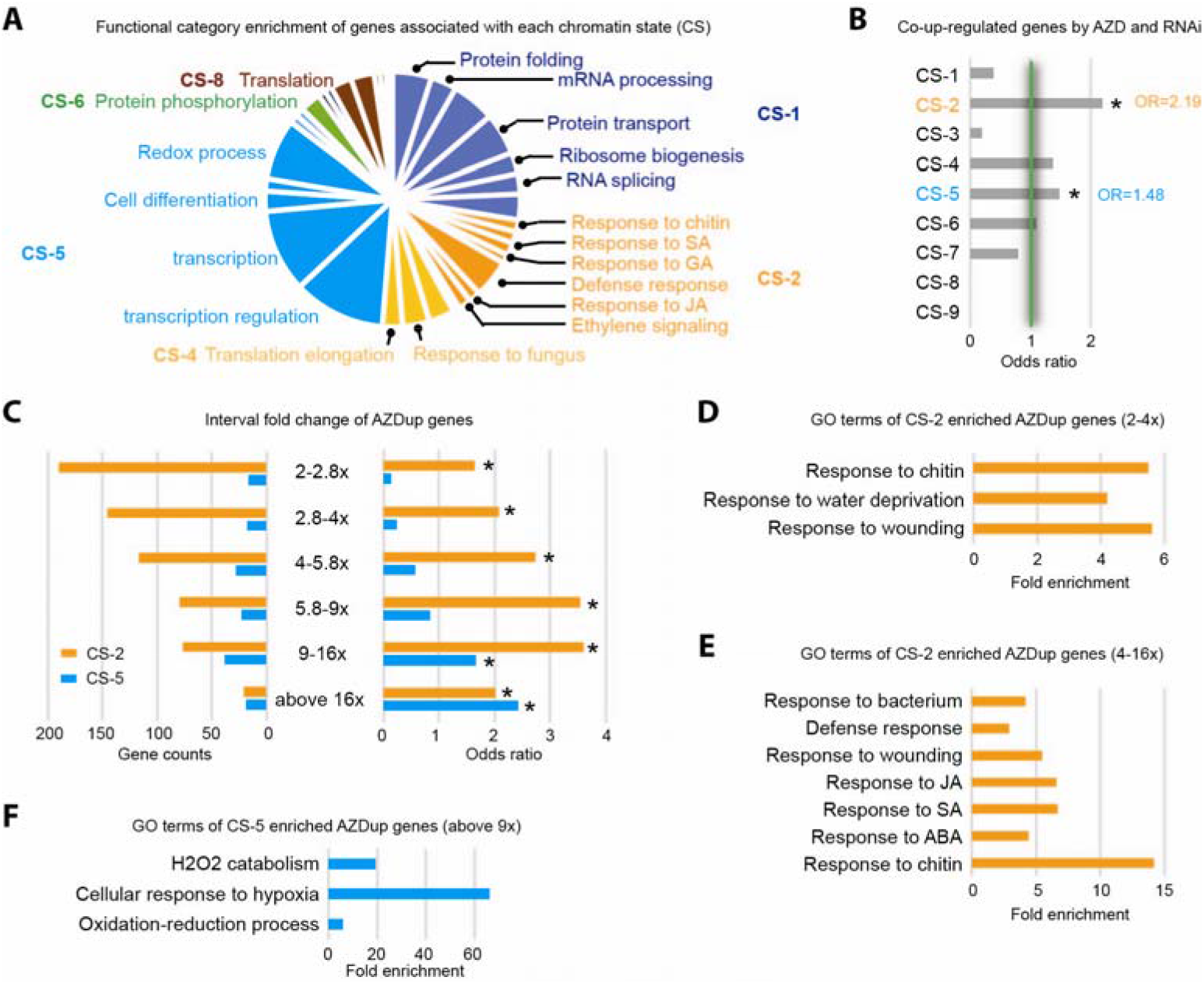
CS-2 and −5 specify the functional groups of TOR-responsive genes. (A) functional category analysis of genes from each chromatin state (CS) (p<0.01, FDR<0.1). Size of pie chart indicates the relative gene number in each category. (B) Enrichment of co-up-regulated genes by both AZD8055 and TOR RNAi in different chromatin states (CS; *, odds ratio>1 and p<0.05). Different chromatin states were defined by Sequeira-Mendes *et al*., 2014. (C) Enrichment and gene counts of different interval fold-change of up-regulated genes by AZD8055 (AZDup) in CS-2 and CS-5 (*, odds ratio>1 and p<0.05). (D-F) Functional category analysis of AZDup CS-2 genes with 2-4 fold, 4-16 fold and CS-5 genes up-regulated at least 9 fold (p<0.01, FDR<0.1).

A list of genes up-regulated (fold change>2.0; p<0.05) upon TOR inhibition by AZD8055 (AZDup, 1583 genes) or RNA interference (TOR-RNAi, 369 genes) was extracted from published datasets (11, 12). 233 genes upregulated in both conditions were significantly enriched at their TSS in the bistable CS-2 and also in the canonical Polycomb CS-5 (Fig. 1B). These findings suggest that the repression of gene expression by TOR is achieved through either CS-2 or CS-5. Furthermore, by distinguishing up-regulated genes depending on their fold change, we observed that the CS-5 enrichment gradually increases with the fold change, which is also the case for CS-2 but not beyond 16-fold change. In addition, more than half of the CS-2 genes change only 2-4-fold (Fig. 1C and Fig. S1). This striking difference could reflect the bistability of CS-2, in which frequent switches between active and silent state may occur, resulting in a low rate of basal transcription, while TOR-repressed genes marked by the monostable silent CS-5 are completely repressed.

Next, using GO analysis, we tested whether AZDup genes marked by either CS-2 or CS-5 are enriched in particular functional categories. AZDup genes marked by CS-2 are enriched in biotic stress and defense mechanisms mediated by phytohormone signaling (Fig. 1D, E), whereas, abiotic stress related genes involved in oxidative stress and hypoxia response are more abundant among AZDup genes with CS-5 features (Fig. 1F). This dichotomy indicates a chromatin state dependent functionalization of TOR repressed genes.

### The chromatin context of TOR-responsive genes is coordinately repressed by CLF and LHP1

Based on the preferential association between TOR repressed genes and CS-2/5, both characterized by the presence of H3K27me3, we hypothesized that a functional link may exist between chromatin factors involved in H3K27me3 deposition and TOR. We tested this hypothesis by collating the TOR repressed gene set with transcript profiles of Polycomb mutants. CURLY LEAF (CLF), a Polycomb group (PcG) protein, is a major H3K27 tri-methylase in plants (13). A total of 84 genes are shared between AZDup and those genes upregulated in the loss-of-function *clf28* and *clf29* mutants (Fig. 2A). These shared genes display a significantly higher association with CS-2 (OR=3.86) at their TSS than the AZDup or the up-regulated genes in either *clf28* or *clf29* (Fig. 2B and Fig. S2). Reinforcing this positive association, the CS-2 enrichment of AZDup decreases when the genes up-regulated in *clf* mutants are excluded (AZDno*clf28/29*; Fig. 2B). Similarly, the CS-5 enrichment is remarkably higher for genes upregulated in both AZDup and *clf* mutants compared to the genes upregulated only in AZDup (AZDno*clf28*/29; Fig. 2B and Fig. S2). Furthermore, the GO analysis of the 84 commonly up-regulated genes between AZDup and *clf* mutants points toward a chromatin state dependent functional categorization. Indeed, commonly up-regulated genes within CS-2 are significantly enriched in biotic and abiotic stress responses, while the ones within CS-5 are specifically enriched in redox processes (Fig. 2C). Then we tested the resistance of the *clf29* mutant to the TOR inhibitor AZD8055. Compared to the wild-type control, *clf29* mutants did not respond to AZD8055 (Fig. 2D), which suggests that growth arrest by TOR inactivation is largely dependent on loss of H3K27me3.

**Fig. 2:**
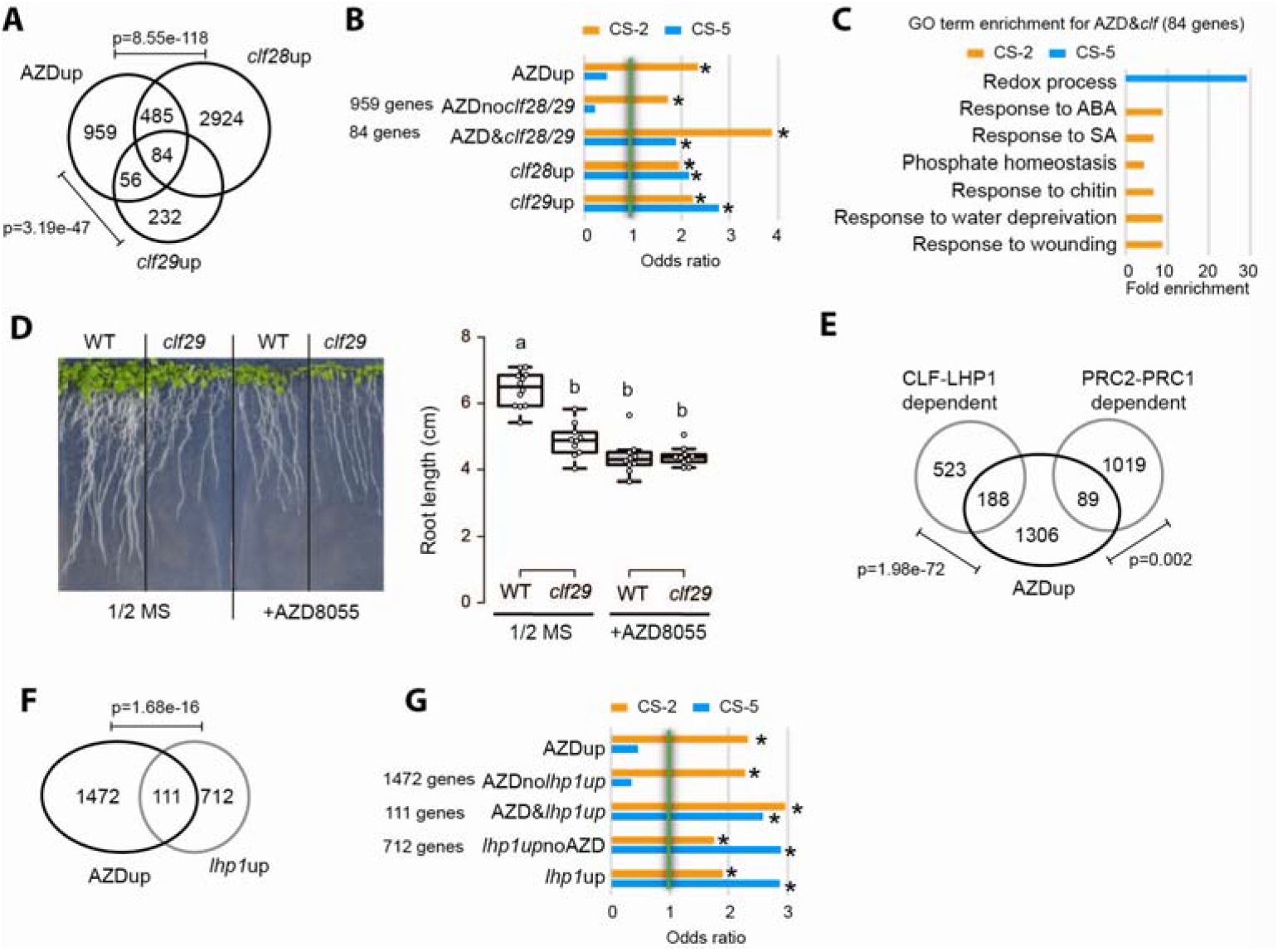
TOR represses both CS-2 and 5 genes via CLF in specific context with LHP1. (A) overlap of up-regulated genes by AZD8055 (AZDup), in *clf28* mutant (*clf28*up, hypergeometric test) and in *clf29* mutant (*clf29*up2x, hypergeometric test). (B) Enrichment of genes from different intersection between AZDup, *clf28*up and *clf29*up presented in (A) (*, OR>1 and p<0.05). (C) Functional category analysis of co-up-regulated genes by AZD8055 and in *clf28/29* enriched in CS-2 and CS-5 (p<0.01, FDR<0.1). (D) WT and *clf29* were germinated and grown on ½ MS medium or ½ MS supplemented with 0.2 μM AZD for 10 days. Bar, 1.5 cm. Root length was determined (n>10, one-way ANOVA, different letters indicated significant difference, p<0.05). (E) overlap of up-regulated genes by AZD8055 (AZDup) and context-dependent CLF regulated genes (hypergeometric test). (F) overlap of up-regulated genes by AZD8055 (AZDup) and in lhp1 mutant (*lhp1*up) (hypergeometric test). (G) Enrichment of genes from different intersection between AZDup and *lhp1*up presented in (F) (*, OR>1 and p<0.05).

PcGs are generally classified into two major complexes named Polycomb Repressive Complexes (PRC1 and PRC2) and CLF, a major PCR2 subunit, selectively collaborates with different PcGs partners to achieve gene repression in distinct developmental programs (14). A first group represents genes commonly up-regulated in *clf* and *lhp1* mutants and may correspond to direct targets of CLF and LHP1 (*i.e*., here after named CLF-LHP1-dependent genes). The second one refers to genes more specifically affected in PRC1 mutants (*i.e*., here after named PRC1-PRC2 dependent genes). Thus, to dissect the relationship between TOR and CLF transcriptional repression, we tested overlaps between these two groups and AZDup genes. A significant overlap was observed between AZDup and CLF-LHP1-dependent genes, while it was not the case with PRC1-PRC2 dependent genes (Fig. 2E). This result suggests that CLF and LHP1 may work concertedly to participate in the repression of TOR specific target genes.

We further investigated the link between TOR and LHP1 by focusing on the 111 genes commonly upregulated by AZD8055 and in the *lhp1* mutant (Fig. 2F). In a manner similar to genes co-upregulated by AZD and *in clf mutants*, these genes showed a significant enrichment in both CS-2 and CS-5 (Fig. 2G and Fig. S3). When we analyzed overlap between LHP1-target genes and genes up-regulated by AZD8055, we found that 445 out of 1583 AZDup genes are indeed targeted by LHP1 (Fig. S4A). CS-2 enrichment of AZDup genes was decreased when LHP1 targets were excluded, whereas genes both upregulated by AZD and targeted by LHP1 present an increased CS-2 enrichment compared to genes *sole* upregulated by AZD or targeted by LHP1 (Fig. S4B). In addition, we noted that the LHP1 targeting also increase the CS-5 enrichment of AZDup genes (Fig. S4B). This observation is reinforced by the fact that when LHP1 direct targets were excluded the association between AZDup and CS-5 dropped dramatically (Fig. S4B). Finally, we investigated whether LHP1 targeting specifies TOR functions using GO analysis. The functional annotation obtained with the CS-2 subset of AZDup genes targeted by LHP1 is mainly linked to biotic and abiotic stress-related functions (Fig. S4C). On the other hand, the functional annotations obtained with the CS-5 subset of AZDup genes targeted by LHP1 appears mainly related to oxidative stress and hypoxia responses (Fig S4D). Together, our results indicate that besides being regulated jointly by CLF and LHP1, TOR repressed targets may further split in distinct functional categories depending on their respective CS-2 or CS-5 chromatin context.

### BRM is important for activating genes associated with CS-2, when TOR is down-regulated in response to biotic stress

TrxG proteins are evolutionarily conserved chromatin-modifying factors that antagonize PcG repression and interactions between PcG/TrxG support a molecular basis for chromatin bistability (7, 15). Among TrxG proteins, the SWItch/Sucrose Non-Fermentable (SWI/SNF)-type protein BRAHMA (BRM) restricts CLF occupancy/activity at many developmental genes (16). We therefore wondered whether the TrxG protein BRM would participate in establishing bistable chromatin at the CS-2 subset of TOR regulated genes. Considering BRM as an activator of gene expression, we observed a significant overlap between AZDup genes and genes down-regulated in *brm1* mutant (Fig. S5) and also found that 57% of AZDup genes are BRM targets (Fig. 3A). Moreover, the CS-2 enrichment among BRM targets was further reinforced when including the overlap with AZDup (Fig. 3B and Fig. S6). Then, our GO analysis demonstrates that AZDup genes which overlap with BRM target genes are mostly biotic stress related, while BRM targets not suppressed by TOR are mostly related to transcriptional regulation (Fig. 3C).

**Fig. 3:**
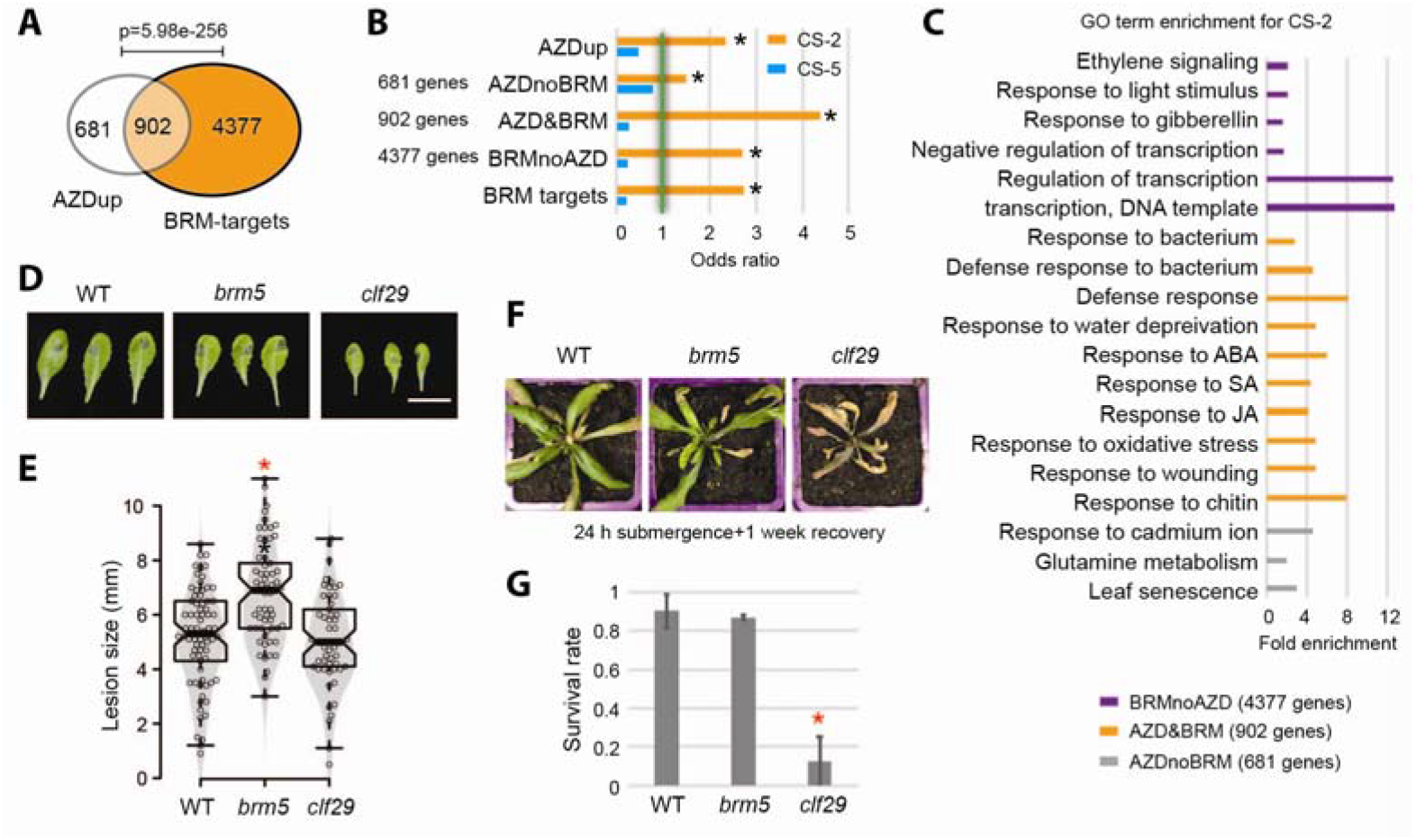
BRM specifically activates CS-2 genes to induce biotic stress response upon TOR inhibition. (A) overlap of up-regulated genes by AZD8055 (AZDup) and BRM target genes (hypergeometric test). (B) Enrichment of genes from different intersection between AZDup and BRM targets presented in (A) (*, OR>1 and p<0.05). (C) Functional category analysis of CS-2 enriched AZDup genes and BRM targets presented in (B) (p<0.01, FDR<0.1). (D) leaf phenotype and (E) lesion size of *brm5* and *clf29* mutants infected by *B. cinerea* compared to col-0 (n>50, central bars of the notched box represent the median, notches indicate 95% confidential interval, *, p<0.05, One-way ANOVA). (F) growth phenotype and (G) survival rate of *brm5* and *clf29* mutants challenged by 24 hours submergence and recovered for another week (n=3, mean±s.d., *, p<0.05, One-way ANOVA).

While keeping in mind the bistability of CS-2, we analyzed the overlap between the TrxG component BRM targets and the PcG component LHP1 targets. A total of 1216 genes are co-targeted by BRM and LHP1, among which 240 genes appeared as AZDup (Fig. S7A). Further reinforcing a functional link between TOR repressed genes and bistable chromatin, these genes show the highest CS-2 enrichment in this study. Furthermore, this result suggests that BRM and LHP1 together act on TOR repressed genes to establish chromatin bistability (Fig. S7B, C). Our GO analysis reveals a preferential enrichment in biotic stress related terms for the CS-2 category, while the Cs-5 was preferentially enriched in GO terms related to oxidative stress and hypoxia. This observation suggests a functional dichotomy between AZDup genes co-targeted by BRM and LHP1 in CS-2 and AZDup genes targeted by LHP1 in CS-5. Notably, the 976 genes co-targeted by BRM and LHP1 but not upregulated upon TOR inhibition preferentially belong to TOR-independent functional categories, *e.g*., salinity response, auxin and gibberellin signaling at CS2; transcription regulation and cell wall modification at CS-5 (Fig. S8).

The proposed functional dichotomy among TOR repressed genes implies that, within CS-2, BRM may specifically activate genes involved in defense mechanism upon TOR inhibition by a biotic stress signal. In agreement, it has been previously shown that over-expression of TOR increased the susceptibility to both bacterial and fungal infections (17). To explore in more details the role of BRM in regulating biotic stress related genes within CS-2, *brm5* mutants were inoculated with the fungal pathogen *Botrytis cinerea*. Compared to the wild-type control (Col-0), *brm5* mutants exhibited an enhanced susceptibility to *B. cinerea* with a significant increase in lesion size while *clf29* did not (Figs 3D, E). Supporting our *in-silico* analyses, opposite sensitivities were observed when mutants were tested for their response to hypoxia caused by submergence. Indeed, while *brm5* behaved similarly as Col0, *clf29* mutants were significantly less resistant to submergence (Figs 3F, G). Together, our results indicate that BRM contributes to the functional specification of TOR responding genes.

### TOR coordinates global H3K27me3 level at different growth and developmental stages

The different correlations highlight a functional link between TOR and H3K27me3 transcriptional repression. We therefore hypothesize that the activation of TOR repressed targets associated with CS-2 and CS-5 may require an active removal of H3K27me3. To test this hypothesis, Arabidopsis were treated for 24 hours with AZD8055 to inhibit TOR. Interestingly, AZD8055 treatment provoked a decrease in the global level of H3K27me3 (Fig. 4A). In a complementary approach, we also explore the impact of a TOR inhibition on the global level of H3K27me3 by means of quantitative immuno-staining on root nuclei. In agreement with the pharmaceutical inhibition of TOR by AZD8055, the level of fluorescence intensity of H3K27me3 was significantly decreased upon inhibition of TOR by estradiol-inducible TOR RNAi with fluorescence levels were intermediate between those of the mock-treated control and the *clf29* mutant (Fig. 4B).

**Fig. 4:**
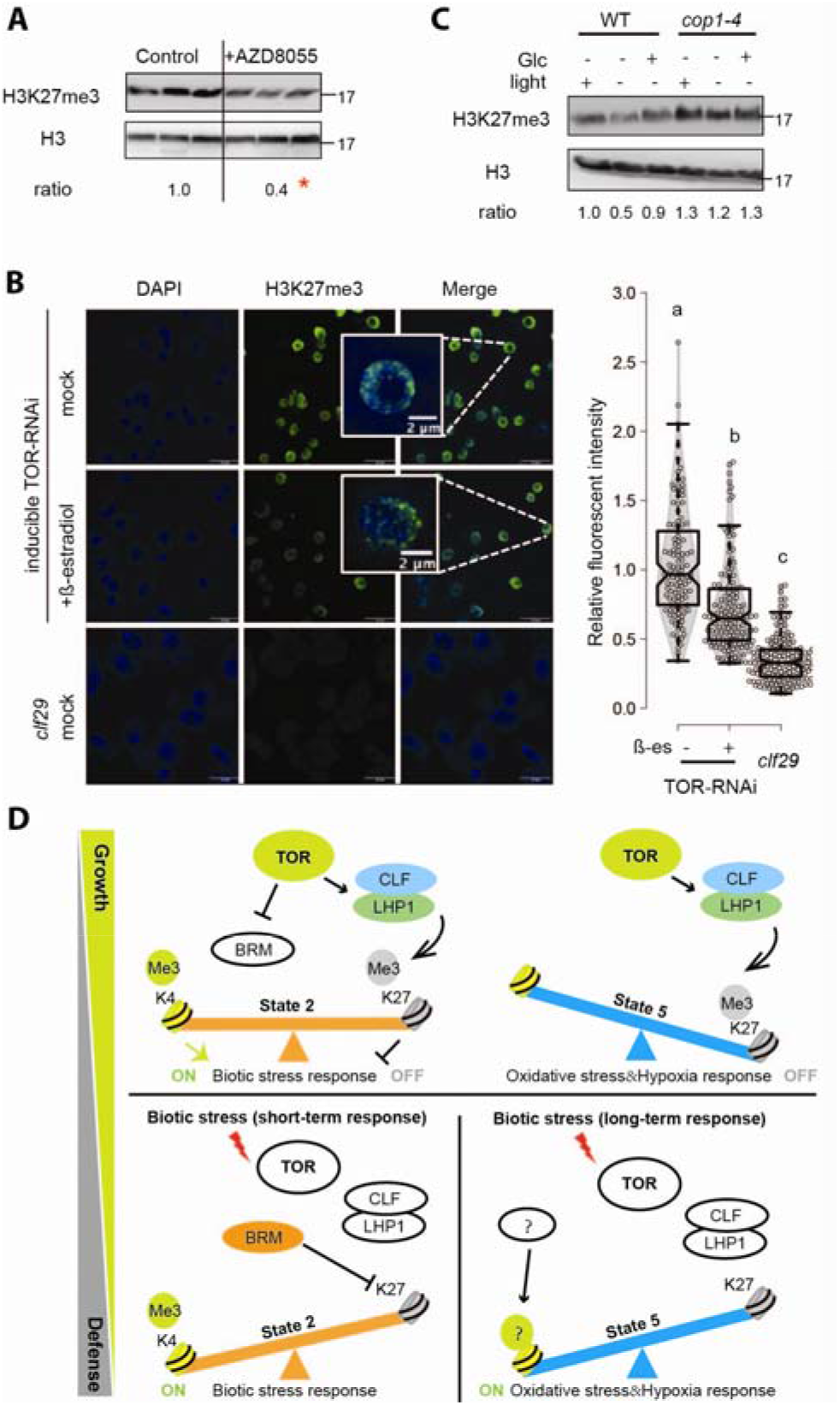
TOR coordinates global H3K27me3 level at different growth and developmental stages. (A) H3K27me3 level was determined in Arabidopsis leaves treated with 1 μM AZD8055 for 24 hours (n=3, t-test, *, p<0.05). (B) Immuno-staining of H3K27me3 in ß-estradiol inducible TOR-RNAi line and *clf29* mutant. Relative fluorescent signal was normalized against DAPI signal (notches indicate 95% confidential interval, letters indicating statistic difference, One-Way ANOVA, p<0.05). (C) WT and *cop1* mutant were grown under different conditions for 4 days. H3K27me3 and H3 level were determined by western blot. (D) Proposed model of TOR function in controlling gene expression to trade-off growth and defense. We propose the following scenario: active TOR up-regulates CLF and LHP1 to deposit the repressive mark H3K27me3 in the bistable chromatin state 2 (CS2) and the Polycomb-associated chromatin state 5 (CS5). TOR inactivation under biotic stress conditions restores the active CS2 state by BRM that removes the repressive epigenetic mark to finetune the rapid biotic stress response. After long-term exposure to biotic stress, plants gradually activate CS5 genes involved in redox process which is more associated with a general stress response.

Both TOR kinase and H3K27me3 are important for photomorphogenesis. In the darkness, TOR activity is repressed by COP1 to allow skotomorphogenesis (*18*). Upon light exposure, genome-wide profiling of H3K27me3 identified a gain in H3K27me3 for more than 2000 genes (*19*). To gain more biological insight, we further tested whether H3K27me3 level is correlated with TOR activity during photomorphogenesis. In agreement with published data (18), light-grown seedlings, synonymous of TOR stimulation, had higher H3K27me3 than dark-grown etiolated seedlings. Similarly, glucose, as a potent TOR activator (3), also provoked an increase in H3K27me3. In the *cop1* mutant where TOR is pre-induced even in the dark (19), H3K27me3 level was also maintained high to allow photomorphogenesis (Fig. 4C). Together, we conclude that TOR can recruit histone-modifying complexes including CLF/LHP1 to promote H3K27 tri-methylation thus ultimately leading to the transcriptome regulation.

## Discussion

In plants, increasing evidence supports that TOR functions as a rheostat to fine-tune the balance between growth and defense (17). However, molecular mechanisms responsible for this balancing operation are still unclear and based on sparse information. Here, we propose a bipartite model in which TOR acts upstream to repress two separate sets of stress related genes by means of two distinct chromatin contexts: *i*) the bistable CS-2 for genes involved in fine-tuning specific stress responses and *ii*) the repressive CS-5 for more general and non-specific stress related genes involved in redox processes and hypoxia responses (Fig. 4D). In agreement with the global effect of TOR inhibition on H3K27me3, we noticed a significant enrichment of CS-2 and CS-5 (*i.e*., two chromatin state characterized notably by the presence of the repressive histone mark H3K27me3) at TOR repressed genes. In addition, we identified significant correlations between TOR-regulated genes and several chromatin factors. LHP1, a plant-specific polycomb subunit, works concertedly with CLF to deposit and spread H3K27me3 and contributes to specify CLF collaborations with different PcGs partners to achieve transcriptional repression in distinct developmental programs (14).

Despite the enrichment in both CS-2 and CS-5 detected among genes up-regulated in *CLF* and *LHP1* loss-of-function mutants, CLF and LHP1 targets were under-represented in CS-2 but preferentially enriched in CS-5 (Fig. S4B and Fig. S9A). In a similar manner, while being significantly enriched in CS-2, genes up-regulated upon TOR inhibition were under-targeted by CLF (Fig. S9B). Considering the bistable nature of CS-2, we believe that these discrepancies may reflect a too transient/dynamic to be captured binding of CLF-LHP1 on CS-2 genes, a labile mechanism proposed as the basis for chromatin bistability (20). Also, these CS-2 genes are not expressed under normal growth conditions due to the presence of H3K27me3. However, H3K4me3 poises them to quickly switch from silent to active transcriptional states in response to TOR inactivation. Such chromatin context may contribute to facilitate a rapid transcriptional response to stress, thus providing an enhanced protection without the costs incurred from the constitutive expression of stress related genes. The coordinate action of CLF and LHP1 seems also to be involved in the deposition of H3K27me3 at CS-5 genes repressed by TOR. CS-2 and CS-5 genes were enriched in distinct functional categories, with CS-5 significantly enriched in genes related to oxidative stress. Interestingly, oxidative stress occurs in response to any kind of environmental cues and antioxidant processes has been proposed to improve the tolerance of plants to various stresses (21). Therefore, the distinctly enriched functional categories we found may reflect different processes of gene expression engagement with CS-2 prevailing for stress response genes and CS-5 for stress tolerance genes.

In an antagonistic manner to CLF-LHP1, we also proposed that the TrxG protein BRM may contribute to regulate CS-2 genes repressed by TOR. This antagonism may occur through its known partner, the H3K27 demethylase REF6 (22). Interestingly, BRM was previously positioned at the nexus of the resource allocation decision between growth and stress responses (23), a positioning which dovetails TOR functions in balancing plant growth and stress responses in response to environmental cues. In addition, BRM is a phosphorylation target of the ABA-SnRK2 relay, which reciprocally regulates the TOR signaling pathway (24, 25). Moreover, BRM protein sequence contains several TOR phosphorylation consensus sites, suggesting that the distinct phosphorylation of BRM by TOR and/or SnRK2 may specify its function (26).

Besides BRM, TOR may regulate H3K27me3 through controlling mRNA translation of several enzymes involved in H3K27 methylation and/or demethylation. La-related protein 1 (LARP1) is a RNA binding protein conserved across eukaryotes and involved in regulating 5’ terminal oligopyrimidine motif (5’TOP) mRNA translation. Recent studies have shown that TOR phosphorylates LARP1 in plants, thus regulating the translation efficiency of 5’TOP-mRNAs that possibly encode proteins involved in chromatin remodeling, e.g. LHP1 (26, 27). In addition, TOR may also regulate the availability of a particular methylation substrate, such as the universal methyl donor S-adenosyl-methionine (SAM) that functions in the methionine-folate cycle. Interestingly, MTHFD1 is a folate cycle enzyme impacting global DNA methylation in plants through SAM availability (28). In animals, mTORC1 senses SAM levels through SAMTOR and MTHFD transcription under tight control of mTORC1 (29, 30). However, it remains unknown whether plant TOR modulates DNA or histone methylation through SAM metabolism in plants. SAMTOR seems to be also a metazoan invention.

In both animals and plants, TOR function is primarily linked to nutrient sensing and translational control. Our findings reveal a novel regulatory role of TOR signaling in chromatin remodeling to modulate specific transcriptional responses. We have identified multiple players involved in this regulation, but additional components are yet to be discovered. In the future, it will be of great importance to dissect this new signaling pathway to understand the role of TOR-dependent epigenetic reprogramming in environmental responses at transitions of developmental stages.

## Methods

### Plant materials and growth condition

*Arabidopsis thaliana* mutant plants, as well as wild-type control plants were the Columbia (Col-0) ecotype. Wild-type plants (N1092) and *brm-5* (N68980) were gained from the European Arabidopsis Stock Centre. *clf-29* (SALK_021003) mutant was described in Wang et al., 2016 (14). *cop1-4* was described in (19). ß-estradiol inducible TOR-RNAi line was described in (3). All seedlings were grown on 1/2MS medium (pH5.7, 0.8% agar) in a long-day climate chamber (16 h light/8 h dark; 80-100 μmol m-2s-1; 22°C day/18°C night; 50% humidity). Chemical inhibitor treatments were performed with media containing 0.2 μM AZD-8055 for 10 days or 1 μM for 24 hours. Submergence was performed with 5 week soil-grown plants for 24 hours and recovered for 1 week in the short-day climate chamber (12 h light/12 h dark).

### Western blot

For immunological detection, total soluble proteins were extracted from 50 mg plant materials with 250 μl 2x Laemmli buffer. Proteins were denatured for 5 min at 95°C and separated on 15% SDS-PAGE. Subsequently, proteins were blotted to PVDF membrane. The primary antibodies anti-H3K27me3 (1:5000, Agrisera, AS16-3193) and anti-H3 (1:5000, Agrisera, AS10-710) were detected using the HRP-conjugated secondary antibody (1:20,000). Full western blot images are provided in Fig. S10.

### Fungi infection

*B. cinerea* was inoculated on 5-week-old soil-grown plants in a short-day chamber (12 h light/12 h dark; 80-100 μmol m-2s-1; 22°C day/18°C night; 50% humidity). For the lesion assay, 5-μL droplets of *B. cinerea* spore suspension were placed directly on the upper surface of the leaf. After inoculation, plants were kept with full humidity to facilitate the infection. Lesions were measured 3 days after the inoculation.

### Chromatin state enrichment analysis

Chromatin states were defined based on Hidden Markov Model (HMM) and previously defined coordinates were extracted (Sequeira-Mendez et al., 2014). The gene sets are given in Supplementary file. Transcription start sites (TSS) and transcription termination sites (TTS) are downloaded from www.arabidopsis.org (TAIR10). An array is formed for each region of interest. GenomicRanges R-package was used to match the genes in each set with a chromatin state (31). The enrichment of chromatin states were calculated by odd ratio (OR) and False Discovery Rate (FDR) adjusted p values were used to define the significance of OR. Statistical comparison of different ORs were performed by Fisher’s exact test and statistical significance threshold is taken as p<0.05. All the gene lists extracted from published datasets and analyzed in this work are provided in Supplementary file.

### Immunostaining and microscopy

Immunostaining was performed as described previously (32) on 7-days-old *in-vitro* grown seedlings of Col-0, TOR-RNAi and *clf-29* treated or not for 24h with 10 μM ß-estradiol. Antibodies used for immunostaining were the anti-H3K27me3 (1:500) and the Alexa fluor-488 dyes-conjugated secondary antibody (1:1000, Life Technologies, A-11008). The H3K27me3 signal intensity for each nucleus was calculated relative to the intensity of DAPI using ImageJ software. Confocal images were acquired with a Zeiss LSM-700 microscope with a 63x/1.01 objective.

## Acknowledgement

We thank D. Janocha (University of Heidelberg) for providing *cop1* mutants, French Agence Nationale de la Recherche—ANR-18-CE13-0019 – ReinitiaTOR to L.R. and 2020 Marie Curie fellowship 885864 TOR in acTIon MSCA-IF-EF-ST to Y.D.

## Author contribution

Y.D. and V.U.U. initiated the project. Y.D., V.U.U., A.B. and C.P. contributed to experiment design and project development. Y.D. coordinated the project and performed experiments. V.U.U. and V.A.S performed bioinformatic analysis. A.B. and G.S. performed immuno-staining. Y.D. and T.H. performed *B. cinerea* infection. Y.D., V.U.U. and A.B. made figures and wrote the manuscript. C.P., T.H. and L.R. revised the manuscript.

## Declaration of interest

The authors declare that the research was conducted in the absence of any commercial or financial relationships that could result in any potential conflict of interest.

## Data availability

All datasets generated for this study are included in the manuscript and/or the Supplementary Files.

## Notes

### Competing Interest Statement

The authors have declared no competing interest.

